# The lysosomal TRPML1 channel promotes breast cancer survival by supporting mitochondrial function and cellular metabolism

**DOI:** 10.1101/2020.09.04.283242

**Authors:** Shekoufeh Almasi, Barry E. Kennedy, Ryan E. Yoast, Scott M. Emrich, Mohamed Trebak, Yassine El Hiani

## Abstract

Triple-negative breast cancer (TNBC) is an aggressive subtype representing approximately 10%-20% of breast cancers and lacking effective therapies. TRPML1, which is a lysosomal Ca2+ release channel upregulated in TNBC, promotes TNBC tumor growth. Here we show a novel crosstalk between lysosomes and mitochondria mediated by TRPML1 in TNBC. TRPML1 is required for the maintenance of mitochondrial function and reactive oxygen species (ROS) homeostasis. TRPML1 knockdown inhibits TNBC mitochondrial respiration, glycolysis and ATP production, leading to reduced proliferation, promotion of cell cycle arrest and apoptosis with enhanced global and mitochondrial ROS. Further, TRPML1 downregulation enhances the cytotoxic effect of Doxorubicin in TNBC cells. Our data reveal a hitherto unknown link between lysosomal TRPML1 channels and mitochondrial metabolism and suggest that TRPML1 inhibition in combination with established chemotherapies could be an effective strategy against TNBC tumors.

## 1. Introduction

A remarkable feature of cancer cells is their ability to grow and proliferate uncontrollably (1). Although the underpinning mechanisms of cancer initiation and progression remain largely obscure, oncogenic alterations leading to enhanced mitochondrial and lysosomal activities are of paramount importance to the survival of cancer cells (2,3).

Cancer cells often exhibit up-regulation of the mitochondrial bioenergetic machinery. Mitochondrial activity supplies cancer cells with ATP to stimulate biomass production and tumor growth but also coordinates distinct metabolic pathways required for cell survival and proliferation (4). Within cells, mitochondria play a key role in shaping both Ca^2+^ and free radical signaling pathways, two fundamental signaling pathways that control cell survival (5,6). Metabolites generated by mitochondria meticulously guide cells through every stage of the cell cycle (7,8). Mitochondrial metabolites also regulate post-translational modifications of proteins (9,10) and modulate the structure and function of chromatin (11–13). Another essential metabolic regulator required for cancer cell survival is the lysosome (14–16). Lysosomes not only execute autophagic degradation of damaged cellular components but also recycle macromolecules to generate building blocks for cellular survival. Lysosomal activity allows cancer cells to cope with starvation and stress conditions while providing a consistent supply of nutrients to fuel malignant growth (17,18). In addition to their involvement in the catabolism and subsequent recycling of biomolecules, lysosomes coordinate a broad range of intracellular signaling pathways, ranging from mTOR, AMPK, Calcineurin, AKT and ERK1/2, all of which contribute to cancer progression (19,20). Importantly, lysosomal function is essential for mitochondrial function and homeostasis, consistent with the role of these two organelles in promoting malignancies (21–23). For instance, the lysosomal biogenesis regulator transcription factor EB (TFEB) is essential for mitochondrial biogenesis, respiratory chain complex activities, and ATP production (24,25), while the endolysosomal Rabs play a central role in mitochondrial oxygen consumption, cytochrome-C release during oxidative stress, and mitophagy (26,27). Disruption of lysosomal acidification diminishes mitochondrial respiration (28) and knockdown of the lysosomal V-ATPase attenuates mitochondrial bioenergetics, induces mitochondrial fission, and increased reactive oxygen species (ROS) production to promote hepatoma cancer cell death (29). Recently, we (30) and others (31,32), demonstrated the role of Transient Receptor Potential Mucolipin 1 (TRPML1 or ML1), a lysosomal Ca^2+^-release channel, in driving breast cancer survival and proliferation. Specifically, we showed that loss of TRPML1 hampers triple-negative breast cancer (TNBC) growth and invasion by modulating mTORC1 activity and lysosomal ATP release to the extracellular space. Here we further show that loss of lysosomal TRPML1 channels impaired the overall cellular metabolism, decreasing mitochondrial respiration and glycolysis flux. This disruption in cellular metabolism resulted in decreased ATP production and cell cycle arrest while also promoting apoptosis and chemosensitivity. Our analysis of OncoGenomics database [GSE22133-GPL5345 (33) and GSE-42568 (34)] complemented our findings and showed that breast cancer patients with low TRPML1 mRNA expression have a better clinical prognosis and higher survival rate. Our results, thus, provide a novel direct functional interface between lysosomes and mitochondria, highlighting TRPML1 as a promising therapeutic target in anticancer therapies.

### 2. Results

#### 2.1 TRPML1 facilitates cell proliferation by promoting apoptotic evasion and cell cycle progression

We recently showed that TRPML1 is specifically upregulated in the triple-negative breast cancer (TNBC) cell lines MDA-MB-231, Hs578T, SUM159PT and HCC38, compared to non-tumorigenic MCF-10A cells and estrogen receptor positive MCF7 cells (30) (see also Fig. 1A). Further, TRPML1 silencing by two distinct shRNAs (KD1 and KD3, Fig. 1B) inhibited the proliferation of TNBC cell lines but not that of the non-tumorigenic MCF-10 A cells (Fig. 1C), suggesting that TRPML1 is vital for the proliferation of TNBC cells. To explore the underlying mechanisms by which TRPML1 knockdown inhibited the proliferation of TNBC cells, we examined whether the knockdown of TRPML1 induced cell cycle arrest. Flow cytometry analysis showed that TRPML1 downregulation markedly increased the percentage of cells in the G0/G1 phase, while the distribution of cells in S and G2 phases decreased (Fig. 1D). To further investigate the molecular mechanism underlying the G1/S phase transition, we synchronized cells at the G0/G1 phase by serum starvation for 24 h, and cells were collected 16 h after re-addition of serum. Western blot analysis was performed to evaluate the expression of key regulators associated with the G1 phase. Consistent with our flow cytometry data, downregulation of TRPML1 significantly decreased the expression of cyclin D1 while increasing the expression of p21 (Fig. 1D). In addition, to test whether knockdown of TRPML1 affected the TNBC cell apoptosis, we evaluated the extent of apoptosis by Annexin V-7ADD staining followed by flow cytometry. The results revealed an elevated level of apoptosis among TRPML1 KD cells (KD1 [20.71%] and KD3 [14.91%]), both of which were significantly higher than that of the empty PLKO vector-transfected control cells [5.41%] (Fig. 1E). Together, our data demonstrate that TRPML1 promotes cell proliferation by promoting G1/S phase transition and facilitating apoptotic evasion.

**Fig.1.**
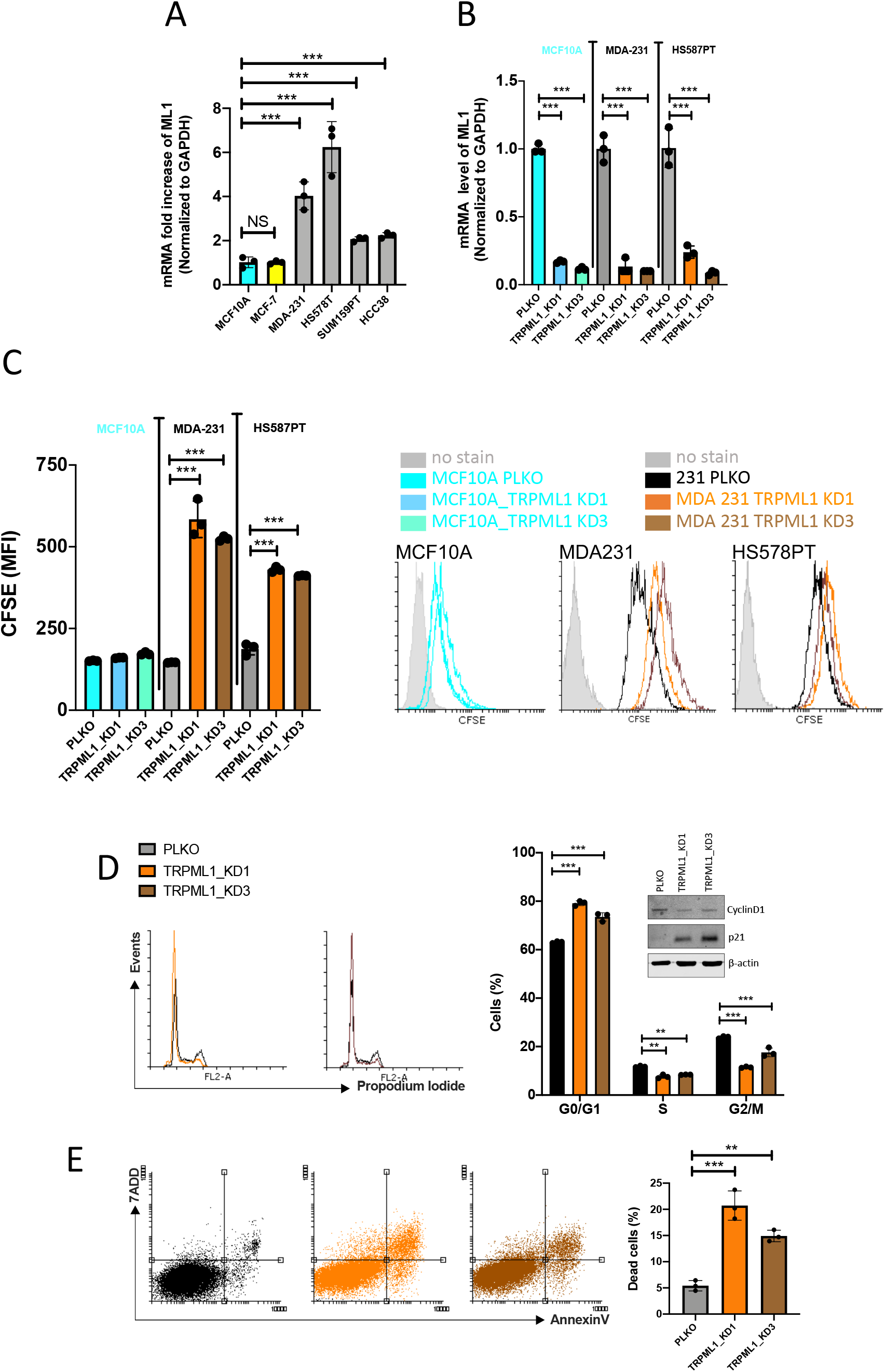
TRPML1 is essential for breast cancer cell survival. A) QPCR analysis of TRPML1 levels in various breast cancer cells compared to non-cancerous control MCF10A cells. TRPML1 is predominant in TNBC MDA-MB231 and HS578PT cells. B) TRPML1 KDs 1 and 3 efficiency tested by QPCR. C) TRPML1 knockdown KDs 1 and 3 significantly decreased proliferation of both MDA-MB231 and HS578PT cells but has no impact on MCF10A control cells (CFSE assay at 72 hours). D) Cell cycle analysis of MDA-MB231 PLKO and TRPML1 KDs stained with propidium iodide and measured 48 hrs after incubation. The histograms denote cell count vs DNA content and the bar graph represent quantitative data. TRPML1 KDs resulted in a statistically significant increase in cells in the G1/0 and decreased cells in the G2/M. E) ML1 KDs promotes apoptosis as reflected by increased cells along annexin V axis and the quantitative analysis in the bar graph. Data represent the mean±SEM (\*\*\**p* < 0.001; \*\**p* < 0.01; *p<0.05; compared to PLKO control cells)

#### 2.2 TRPML1 knockdown alters cell metabolism of triple-negative breast cancer MDA-MB231 cells

One hallmark by which cancers meet their biosynthetic demands is through global reprogramming of cellular metabolism (35,36). To investigate the link between TRPML1 and metabolism, we performed targeted mass spectrometry-based metabolomics on MDA-231 and MCF10A cells with and without TRPML1 knockdown. In line with a non-essential role of TRPML1 in non-cancerous MCF10A cells, we observed minimal changes in the levels of 145 metabolites (Fig. 2A). Conversely, depletion of TRPML1 in MDA-MB231 cells resulted in global metabolic alterations as compared to control (Fig. 2B). This suggested to us that altered metabolism was potentially responsible for the reduced rate of cell proliferation by TRPML1 knockdown in cancerous cells. To identify metabolic pathways that could contribute to the reduced cell proliferation, an enrichment analysis was performed on metabolites that were increased or decreased by a minimal of 1.5-fold in MDA-MB231 cells with TRPML1 knockdown as compared to control (Fig. 2C). Enrichment analysis identified several critical metabolic pathways altered by changes in TRPML1 expression levels in MDA-MB231 cells including amino acid metabolism, nucleic acid metabolism, and mitochondrial metabolism. Specifically, levels of the amino acids L-proline and L-glutamic acid were significantly enhanced by TRPML1 knockdown in MDA-MB231 (Fig. 2D). Furthermore, L-arginine levels were significantly decreased, while L-glutamine and L-histidine were not significantly altered upon TRPML1 knockdown in MDA-MB231 cells as compared to control (Fig. 2D). Another top hit identified by enrichment analysis was nucleic acid metabolism, nucleic acid intermediates xanthosine, guanosine triphosphate, and inosine were significantly increased whereas deoxy GTP was significantly decreased by TRPML1 knockdown in MDA-MB231 cells as compared to control (Fig. 2E). Changes in amino acids and nucleic acid metabolism are well known to occur during changes in cell proliferation, and the exact contributions by these pathways in TRPML1-dependent cell proliferation require further exploration. The metabolic pathway that we focused the rest of our study on was mitochondrial metabolism. Consistent with mitochondrial dysfunction upon TRPML1 knockdown, altered metabolites were commonly grouped in citric acid cycle, oxidation of branched chain fatty acids, malate-aspartate shuttle, and Warburg effect (Fig. 2C). Specifically, steady-state levels of TCA cycle intermediates, isocitrate acid and succininic acid were significantly increased by knockdown of TRPML1 in MDA-MB231 but not in MCF10A (similar trend was observed for L-malic acid while no significant decrease in oxoglutaric acid was observed) (Fig. 2F). These results would be in line with a disruption in the TCA cycle, which would partially explain how TRPML1 knockdown would slow cell proliferation of cancerous cells.

**Fig.2.**
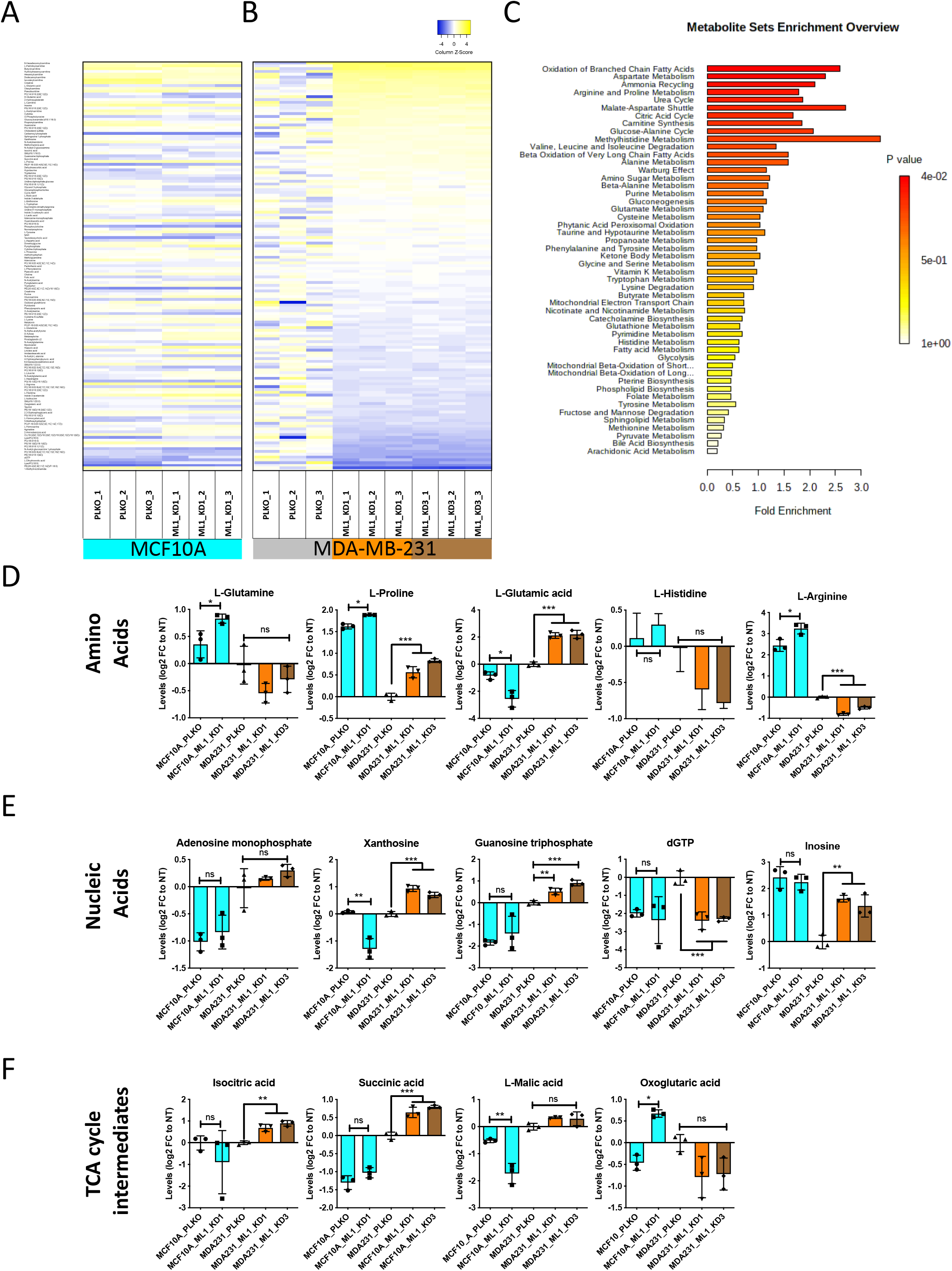
ML1 control cell metabolomics. We performed targeted mass spectrometry-based metabolomics on MDA-MB231 and MCF10A cells in the context of TRPML1 KD and observed changes in serval metabolites caused by TRPML1 knockdown (ML1_KD) in MDA-MB231 cells, while no obvious changes can be seen in MCF10A cells (A). To identify metabolic pathways that could contribute to the enhanced cytotoxicity, an enrichment analysis was performed on metabolites that were significantly changed by TRPML1 knockdown as compared to PLKO (A and B). These results would be in line with a block in the TCA cycle. Additionally, alterations in amino acids [C] (i.e. L-proline, L-glutamic acid, L-histidine and L-arginine) and nucleic acids [D] (i.e. xanthosine, dGTP, inosine) are likely linked to the decreased proliferation and block in cell cycle in MDA-MB231 cells with knock down of TRPML1. In line with mitochondria dysfunction, altered metabolites were commonly grouped in citric acid cycle, oxidation of branched chain fatty acids, malate-aspartate shuttle, and Warburg effect. Furthermore, steady-state levels of TCA cycle intermediates, isocitrate acid and succininic acid were significantly increased by knock down of TRPML1 in MDA 231 but not MCF10A (similar trend for L-malic acid and a non-significant decrease in oxoglutaric acid) (E). Data represent the mean±SEM (\*\*\**p* <0.001; \*\**p* <0.01; *p<0.05; compared to PLKO control cells).

#### 2.3 TRPML1 positively regulates mitochondrial respiration, ATP production and glycolytic flux

Because mitochondrial function is vital in the synthesis of adenosine triphosphate (ATP), which controls a plethora of cellular activities, such as cell cycle progression, cell division and survival (37), we examined whether TRPML1-mediated TNBC survival involves regulating mitochondrial bioenergetics. To expand our analysis of mitochondrial function in response to TRPML1 knockdown, we performed mitochondrial stress tests using seahorse XF24 extracellular flux analyser (38). We first measured mitochondrial oxygen consumption rates (OCRs) during the subsequent addition of mitochondrial inhibitors: oligomycin, FCCP and rotenone + Antimycin [Rot]), in both control PLKO and TRPML1 KDs of MDA-MB231 TNBC and non-cancerous MCF10A control cells. As shown in Fig. 3A, knockdown of TRPML1 drastically decreased OCR of MDA-MB231 cells but had no impact on OCR of the non-cancerous MCF10A control cells (Fig. 3A). In fact, knockdown of TRPML1 in MDA-MB231 reduces the OCR levels to those of non-cancerous MCF10A control cells (Fig. 3A). Furthermore, TRPML1 knockdown reduced both basal respiration (Fig. 3B) and maximal respiratory capacity (OCR after subsequent additions of oligomycin and FCCP) (Fig. 3C) of MDA-MB231, while it does not affect such aspects of MCF10A control cells, suggesting that TRPML1 is an essential contributor to the oxidative phosphorylation of MDA-MB231 cancer cells. On the other hand, because glycolysis plays a significant role in cancer cell proliferation (39,40), we examined the influence of TRPML1 on glycolysis using seahorse glycolysis stress test. Interestingly, TRPML1 knockdown significantly reduced the extracellular acidification rate (ECAR) (Fig. 3C) suggesting decreased glycolytic flux (Fig. 3D) and glycolytic capacity (ECAR after oligomycin addition) (Fig. 3E) in MDA-MB231 cells with no impact on the non-cancerous MCF10A cells. Importantly, ATP production (inferred from OCR after oligomycin addition) was impaired in TRPML1 KD MDA-MB231 cells but not in MCF10A cells (Fig. 3G). Assessment of the relative utilization of mitochondrial respiration versus glycolysis under basal conditions confirmed that knockdown of TRPML1 reduced both mitochondria-mediated oxidative phosphorylation and the glycolytic potential of TNBC cells, thus suppressing the overall metabolic rate of cancerous cells (Fig. 3H). These results suggest that TRPML1 contributes to TNBC malignancy by promoting the metabolic activity and mitochondrial bioenergetics of cancerous cells.

**Fig.3.**
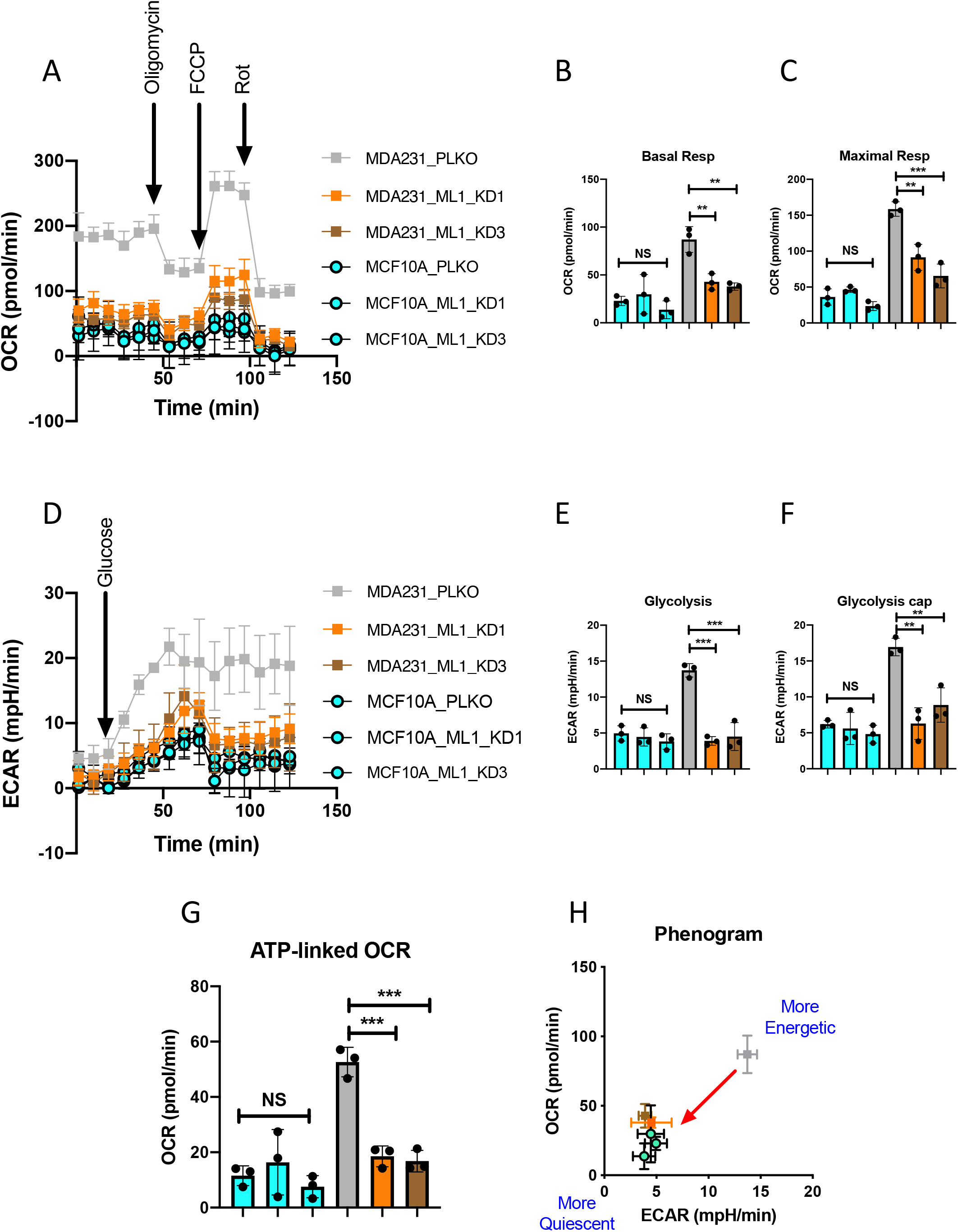
TRPML1 is required for mitochondrial function and bioenergetics. A) TRPML1 KD markedly reduces OCR of MDA-MB231 to levels of MCF10A control cells, while it has no impact on MCF10A control cells. TRPML1 knockdown significantly decreased MDA-MB231 basal respiration (B) and maximal respiration (C). D) TRPML1 KD reduces MDA-MB231 ECAR but has no impact on MCF10 control cells. TRPML1 downregulation inhibits mitochondrial glycolysis (F), and glycolysis capacity (G), while it has no impact on MCF10 control cells. G) TRPML1 KD markedly reduces MDA-MB231 ATP production but has no impact on MCF10 control cells. H) TRPML1 downregulation turndown MDA231 from highly to low metabolic rate. Data represent the mean±SEM (\*\*\**p* <0.001; \*\**p* <0.01; *p<0.05; compared to PLKO control cells).

#### 2.4 TRPML1 knockdown enhances ROS production without affecting mitochondrial membrane potential (ΔΨm)

We then investigated the potential role of TRPML1 in maintaining low homeostatic levels of reactive oxygen species (ROS) within cancer cells necessary for survival and apoptosis evasion. We analyzed the basal levels of whole-cell ROS (using H2DCFDA) and mitochondrial ROS (using MitoSOX Red) of PLKO control and TRPML1 KD of MDA-MB231 TNBC and non-cancerous MCF10A control cells (41). In the non-cancerous MCF10A cells, our results showed no significant difference of ROS (both total and mitochondrial) levels between PLKO and TRRPML1 KD (Fig. 3A and 3B). However, in MDA-MB-231, TRPML1 knockdown resulted in a significant increase in both the total (Fig. 4A) and mitochondrial (Fig. 4B) ROS levels. Our data demonstrates the important role of TRPML1 in regulating ROS production in TNBC, exerting protective / anti-apoptotic effects to support cancer cell growth. Considering that ROS production is positively correlated with mitochondrial membrane potential (ΔΨm) (42,43), we next examined whether TRPML1 influences ΔΨm. Control PLKO and TRPML1 KD of both MDA-MB231 TNBC and non-cancerous MCF10A control cells were pre-treated with the tetramethylrhodamine ethyl ester (TMRE) and then analysed using flow cytometry. Intriguingly, no difference of ΔΨm levels was detected between these cell groups, suggesting that TRPML1-mediated ROS regulation is independent of ΔΨm in these cells (Fig. 4C).

**Fig.4.**
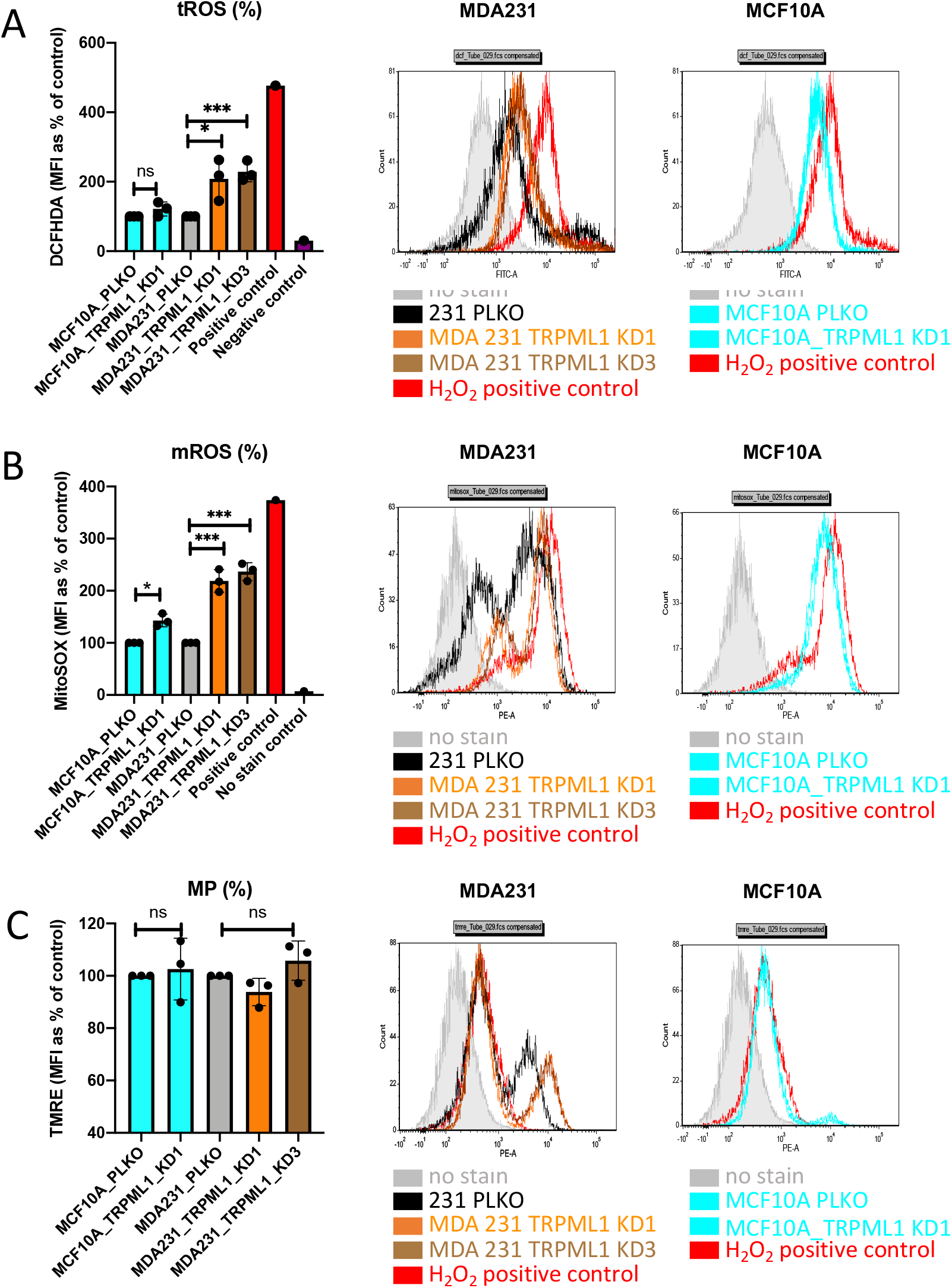
TRPML1 control ROS production but has no impact on mitochondrial membrane potential. TRPML1 KD markedly promotes the production of total ROS (A) mitochondrial ROS (B) in MDA-MB231, but not in MCF10A control cells. C) TRPML1 KD did not affect mitochondrial membrane potential either in MDA-MB231 or MCF10A. Data represent the mean ± SEM (*\*\*p* < 0.001; *\*\*p* < 0.01; *p<0.05; compared to PLKO control cells)

**Fig.5.**
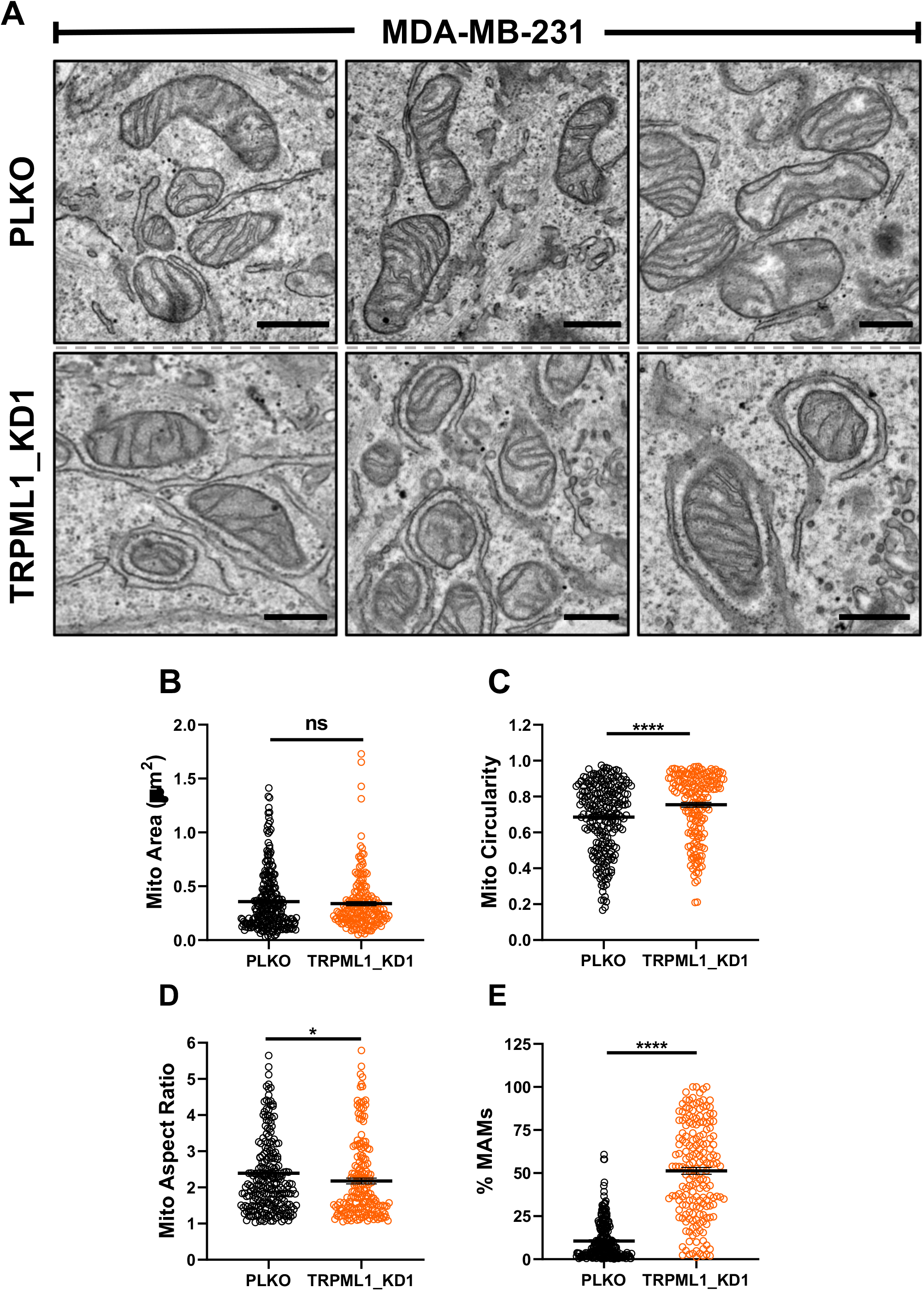
TRPML1 down-regulation alters mitochondrial shape and mitochondria-associated membranes. (A) Three representative electron micrographs displaying mitochondria from PLKO and TRPML1_KD1 transduced MDA-MB231 cells (Scale bar = 0.5 μm). (B-E) Quantification of mitochondrial cross-sectional area (p=0.8670), circularity (p<0.0001), aspect ratio (p=0.0292), and percentage of MAMs (p<0.0001), respectively. All datapoints represent individual mitochondrion (PLKO: n=245; shTRPML1: n=186). For each condition at least 15 electron micrographs were analyzed. Statistical significance was determined using the Mann-Whitney test (ns, p>0.05; *p<0.05; ****p<0.0001).

#### 2.5 TRPML1 knockdown alters mitochondrial shape and mitochondria-associated membranes (MAMs)

We used electron transmission microscopy to analyze the ultrastructural properties of mitochondria in control PLKO-transfected and TRPML1 KD MDA-MB231 cells. While the overall number of mitochondria and cristae structure did not significantly change in TRPML1 KD MDA-MB231 cells compared to control, mitochondria of TRPML1 KD cells were more spherical (Figure 5). Intriguingly, the proportions of endoplasmic reticulum membranes closely associated with mitochondria (MAMs for mitochondria-associate membranes) was significantly higher is TRPML1 KD MDA-MB231 cells (Figure 5).

#### 2. 6 TRPML1 downregulation enhances the cytotoxic effect of Doxorubicin in triplenegative breast cancer MDA-MB231 cells

To assess the therapeutic relevance of our findings, we investigated whether TRPML1 silencing can influence MDA-MB231 cells response to doxorubicin, a widely used chemotherapeutic agent for early and advanced breast cancer therapy (44,45). Interestingly, our results show that TRPML1 downregulation enhanced chemo-sensitivity to doxorubicin, further inhibiting cell viability upon exposure to doxorubicin (Fig. 6A). For instance, 120 nM of doxorubicin at 24 hours has almost no effect on PLKO MDA-MB231 cell (11.97 % of decreased viability), while the same dose of doxorubicin induces a significant reduction in cell viability of TRPML1 KD MDA-MB231 cells (32% and 49.23% of decreased viability in TRPML1 KD1 and KD3, respectively). Similar results were obtained at 48- and 72-hours (Fig. 6B, C). Our results illustrate the therapeutic potential of TRPML1 inhibition in potentiating the anti-tumoral effects of doxorubicin. Consequently, applying TRPML1 inhibitors with conventional chemotherapeutic agents, should be further explored in future antineoplastic therapies.

**Fig.6.**
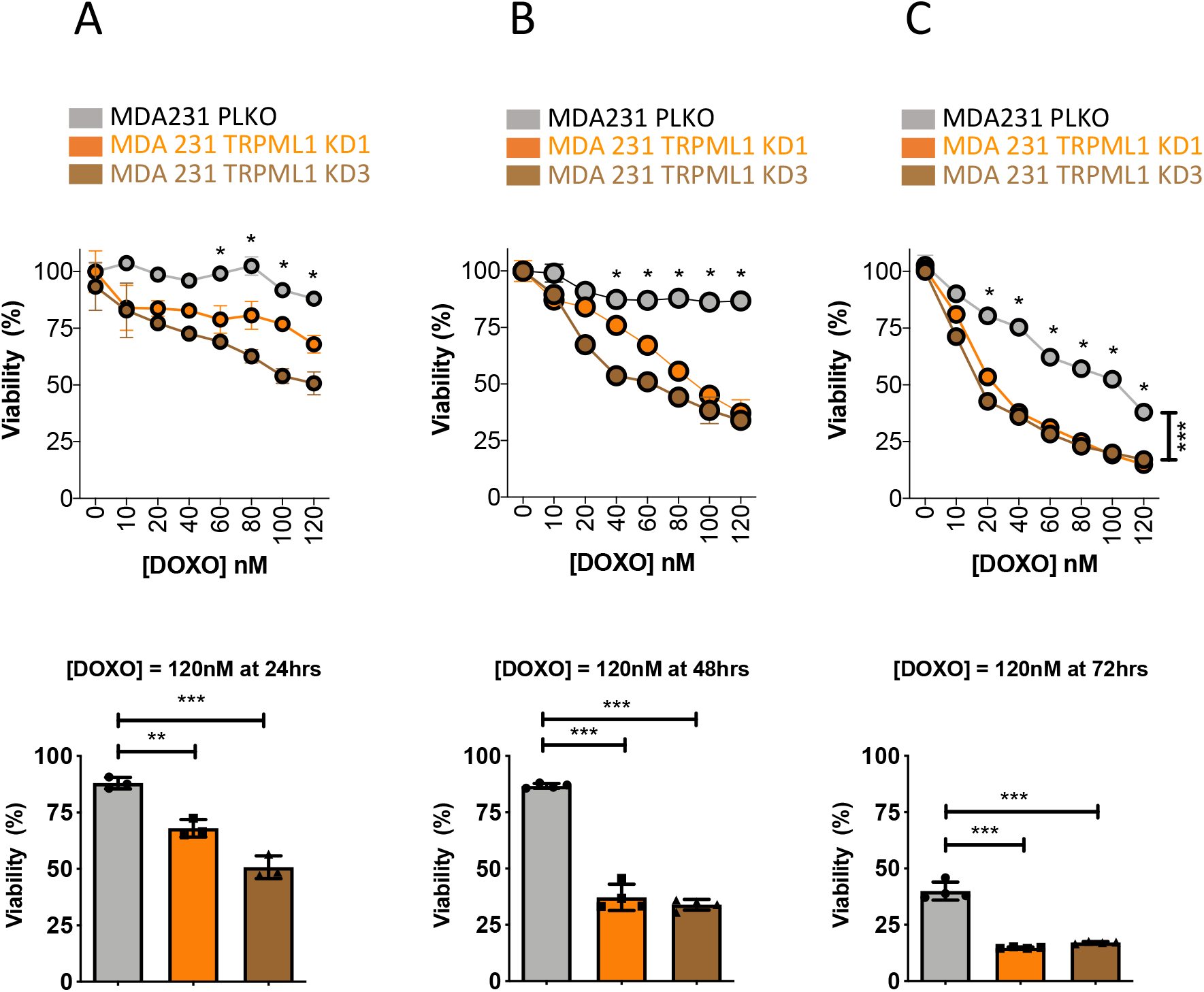
TRPML1 down-regulation enhances the efficacy of doxorubicin (DOXO) in a dosedependent manner in MDA-MB231 TNBC. Dose response curve of DOXO at several time points 24h (A), 48h (B) and 72h (C) in PLKO (gray) and ML1 KDs 1 (orange) and 2 (brown) of PLKO and TRPML1 depleted MDA-MB231 cells. Data represent the mean ± SEM (\*\*\**p* < 0.001; *\*\*p* < 0.01; compared to PLKO control cells)

### 3. Discussion

The lysosomes, which are conventionally known for processing and recycling macromolecules, have garnered increasing attention and gradually emerged as key regulators of autophagy, nutrient sensing, and cellular metabolism. Here, we unravel a novel role for the lysosomal Ca^2+^ release channel TRPML1 channel in breast cancer metabolism. We show that TRPML1 is required for efficient mitochondrial function and survival of triple-negative breast cancer cells. We also show that TRPML1 inhibition sensitizes cancer cells to doxorubicin chemotherapy. The loss of TRPML1 induces a series of metabolic rewiring that diminishes mitochondrial function and bioenergetics, which ultimately leads to enhanced ROS production, abnormal cell cycle progression, apoptotic induction, and potentiation of chemosensitivity. These findings are consistent with previous studies that showed the importance of lysosomes in metabolic reprograming of cancer cells (46,47).

Our study reveals a strong functional link between the lysosomal TRPML1 channels and mitochondria in cancer survival. A recent study identified sites of close contact between lysosomes and mitochondria and showed that TRPML1 mediates lysosomal Ca^2+^ release, which propagates into mitochondria through these mitochondria–lysosome contact sites (48). Therefore, TRPML1-mediated Ca^2+^ transfer to mitochondria is likely fueling bioenergetics in a manner similar and complementary to ER-mitochondria Ca^2+^ transfer (49). This is consistent with our results showing that the mitochondrial membrane potential is not altered when TRPML1 is downregulated. The enhanced mitochondria-associated ER membranes (MAMs) in TRPML1 knockdown cells is likely a compensatory mechanism in response to decrease lysosomal Ca^2+^ transfer to mitochondria. Clearly, further organellar-specific Ca^2+^ measurements studies are needed to understand the relative contribution of TRPML1-mediated Ca^2+^ transfer to mitochondria in cellular metabolism.

Recent data from our group showed that TRPML1 channels are upregulated in TNBCs and play a role in promoting TNBC progression through mTORC1 activity and release of lysosomal ATP into the extracellular space (30). TRPML1 was shown to be a ROS sensor that is required for clearing excess ROS within the cells and preventing oxidative stress (50). These findings are consistent with our data herein showing an increase in ROS production when TRPML1 is knocked down in TNBC. Therefore, our data is consistent with a role for TRPML1 in maintaining the functional integrity of the mitochondria and supplying mitochondrial bioenergetics, to facilitate TNBC cell survival and disease progression. First, we show that loss of TRPML1 causes disturbances in overall cell metabolism, altering the biosynthesis of various amino acids and nucleic acids, both of which are indispensable components for cancer growth and progression (51,52). For example, amino acids L-proline, L-glutamic acid, and L-arginine and nucleic acids xanthosine, guanosine triphosphate, and inosine were significantly impaired upon TRPML1 downregulation in TNBC cells. Changes in amino acids and nucleic acid metabolism often accompany changes in cell proliferation, and further studies will be necessary to determine the exact contributions by these metabolites in TRPML1-dependent cell proliferation. Second, TRPML1 inhibition suppressed the metabolic activity of MDA-MB231 cells. The metabolic suppression was manifested in reduced mitochondrial OCR, glycolysis, and ATP production. In line with mitochondrial dysfunction following TRPML1 knockdown, various TCA cycle intermediates were also found altered in TRPML1-depleted TNBC cells. For example, isocitrate acid and succininic acid were significantly increased in TRPML1 depleted compared to PLKO MDA-MB231 cells. Third, TRPML1 downregulation significantly induced total and mitochondrial ROS production. ROS are by-products of biological reactions with a double-edged-sword effect in oncogenesis (53). Moderate elevation of ROS facilitates the activation of a large spectrum of intracellular signaling which ultimately leads to tumour progression (54). However, excessive ROS induces severe damages to vital cellular structures, such as the mitochondria and the nucleus, ultimately instigating apoptosis (54,55).

One important observation from our study is the selective absence of deleterious effects associated with TRPML1 downregulation in non-cancerous cells. Indeed, TRPML1 knockdown did not exert any major impacts on MCF10A cell metabolism, mitochondrial function/bioenergetics, ROS production or proliferation. This indicates that pharmacological agents inhibiting TRPML1 are expected to selectively kill cancerous cells. We show that TRPML1 downregulation significantly enhances the sensitivity of TNBC cells to doxorubicin. Given that doxorubicin exerts its cytotoxic effect on tumor cells mainly through the generation of ROS (56), we speculate that increased efficacy of doxorubicin in TRPML1-depleted cells may be facilitated by increased ROS as a result of TRPML1 knockdown.

In summary, we discovered an essential functional connection between the lysosomes and mitochondria, where TRPML1 maintains cellular metabolism, enhances mitochondrial function and bioenergetics, and limits ROS generation in the pathophysiology of TNBC. Importantly, the role of TRPML1 is specific to cancer cells as inhibition of TRPML1 do not affect non-cancerous cells. Thus, the differential regulation of TRPML1 in oncogenic states opens new avenues for pharmacological interventions. Future research will focus on the identification of signaling pathways involved in TRPML1-mediated cellular metabolism to oncogenesis.

## 4. Material and methods

### 4.1 Cell culture

Human immortalized breast normal (MCF10A: ATCC-CRL-10317) and cancer cell lines (MCF7: ATCC-HTB-22, MDA-MB-231: ATCC-HTB-26, HS578t: ATCC-HTB-126, SUM159, HCC38: ATCC-CRL-2314) were purchased from ATCC and generously given by Dr. Marcato. Cells were cultured in the specific media, supplemented with 10% heat-inactivated fetal bovine serum (FBS; Gibco, Life Technologies, 16000036), and 20 μg/ml penicillin/streptomycin antibiotic (Gibco; Life Technologies, 15070063) at 37°C and 5% CO2 incubator.

### 4.2 Generation of stable gene-knockdown cell lines

Gene-specific shRNA sequences cloned into the TRC cloning vector (pLKO.1 puro plasmid) were purchased from Dharmacon. Lentiviral particles were produced in HEK-293 cells according to the protocol of the 3rd generation lentiviral packaging system. Cells cultured in 10 cm-plates reaching sixty percent confluency were co-transfected with PPAX2 (6 μg), MD2G (3 μg) and pLKO-LV-gene-specific (6 μg) plasmids in the presence of PEI transfection reagent (Sigma). The generated lentiviral particles were collected 24 hours post-transfection, filtered (Millex-GS; 0.22 μm sterile filter) and stored at −80 °C. Cancer cells were grown in 6-well plates for 24 hours and transduced with 500 μL of lentivirus aliquot diluted in cell-specific medium in the presence of 8 μg/mL of Sequebrene (Sigma) and incubated at 37°C and 5% CO2 for 72 hrs. Transduced cells were selected using 1 μg/mL puromycin for 48 hrs. Lentiviral knockdown efficiency was assessed using RT-qPCR.

### 4.3 RNA isolation and RT-qPCR

Whole mRNA was extracted using Invitrogen RNA Purification kit according to the manufacturer’s protocol. 2 μg of purified RNA was used in the synthesis of complementary DNA (cDNA) based on Super Script® II First-Strand Synthesis System (Invitrogen). Quantitative PCR was performed to examine the expression of genes of interest according to BioRad protocol using CFX96 touch real-time PCR instrument (BioRad). Gene expression data was analyzed using Livak and Schmittgen’s 2-ΔΔCT method and normalized to the 3-phosphate dehydrogenase (GAPDH) reference gene. Primers’ sequences for GAPDH (CTGAAGAGCTGCTTCACCAA/ATGGT GCTGTCCTTGACAAC) and for TRPML1 (TCTTCCAGCACGGAGACAAC/ GCCACATGAACCCCACAAAC).

### 4.4 Western

1x RIPA buffer (20 mM Tris-HCl (pH 7.5), 150 mM NaCl, 1 mM Na2-EDTA, 1 mM EGTA, 1% NP-40, 1% sodium deoxycholate, 2.5 mM sodium pyrophosphate, 1 mM β-glycerophosphate, 1 mM Na_3_ VO4, 1 μg/ml leupeptin) was used to extract protein from cells. Isolated protein samples were quantified according to the BCA assay protocol (Thermofisher Scientific). Twenty μg of each protein sample was separated using SDS-gel electrophoresis and transferred onto a 0.45-μm PVDF membrane (BioRad). Membranes were incubated with blocking buffer (5% milk powder dissolved in 1x TBST) for 1 hour at room temperature and kept in specific primary antibodies overnight at 4°C (β-Actin: 3700s, p21 Waf1/CIP1: 2947s, Cyclin D2: 3741s). The following day, membranes were washed with 1x PBS and incubated with proper fluorescent secondary antibody (Goat-Anti-Mouse, Goat-Anti-Rabbit; Mandel Scientific) for 1 hour at room temperature. After a wash, blots were scanned using the Li-Cor Odyssey 9120 infrared imager.

### 4.5 Cell proliferation assay

Cells in suspension were stained with 100 μL of 2.5μM CFSE (Carboxyfluorescein succinimidyl ester; Sigma) for 10 min in the dark at 37°C. Stained cells were cultured in 12-well plates at 37°C and 5% CO2 for 4 days. On the day of the experiment, cells were washed and suspended in FACS buffer (1% FBS and 1% 0.5 M EDTA in 1x PBS) and fluorescent intensity was determine using BD FACSCaliburtm (Spectron Corporation) at a wavelength of 488nm. The reduced fluorescent intensity indicates higher proliferation rate. Data analysis was performed by Flowing software 2.5.1.

### 4.6 Annexin v/7-AAD binding assay

Annexin V/7-AAD binding assay was performed to quantify the percentage of apoptotic and necrotic cells. Cells were grown in 6-well plates at 37°C and 5% CO2 for 72 hours. On the day of the experiment, cells were washed with 1x Annexin buffer (0.1M HEPES/NaOH (pH 7.4) 1.4 M NaCl, 25 mM CaCl2) and stained with 12.5 μg/mL AnnexinV-fluorescein isothiocyanate (AnnexinV, Alexa Fluor 488; Invitrogen) and 20 μg/mL 7-AAD solution (7-amino-actinomycin D, Biolegend, 650 nm) for 15 min at room temperature in dark. The stained cells were then re-suspended in 1 mL of 1x Annexin buffer and the mean fluorescent intensity was quantified using the BD FACSCaliburtm.

### 4.7 Cell cycle analysis

To determine the percentage of cells in distinct phases of cell cycle, Propidium Iodide (PI) staining was performed. Briefly, 10^5^ cells were seeded in 6-well plates and synchronized by FBS starvation for 24 hours. Next day, culture medium was first replaced with complete growth medium containing 10% FBS followed by incubation at 37°C and 5% CO2 for 72 hours. The day before the experiment, cells were washed with 1x PBS and fixed with ice-cold 70% ethanol overnight at 4°C. On the day of the experiment, cells were washed with 1x PBS twice and stained with 1 mL of PI solution (100 μg/mL PI and 50 μg/mL RNase A in 1x PBS) for 1 hour at room temperature in dark. Samples were read using BD FACSCaliburtm (Spectron Corporation) at 488-nm wavelength with data being processed by the Flowing 2.5.1 software.

### 4.8 Global metabolomics

Metabolites were extracted from cells by scraping in cold (−20°C) 80% methanol. Samples were centrifuged at 13,000xg for 5min and a 25μL aliquot of supernatant was added to 225μL of hydrophilic interaction liquid chromatography (HILIC) loading buffer containing 95% acetonitrile, 2mM ammonium hydroxide, and 2mM ammonium acetate then centrifuged again at 13,000xg for 5 min. Triplicate, 50μL injections of the supernatant were loaded on an Acquity UPLC BEH Amide, 1.7um particle size, 2.1×100mm column (Waters #186004801). Multiple reaction monitoring (MRM) was performed using a Sciex 5500 QTRAP using a previously described acquisition method (57,58). Peak heights for individual metabolites were extracted using Skyline software (MacCoss Lab).

### 4. 9 Extracellular flux analysis

Seahorse XF24 Extracellular Flux Analyzer (Seahorse Bioscience, Billerica, MA, USA) was used to assess mitochondrial function as previously described. Briefly, 10^5^ cancer cells were seeded into the XF24 cell culture microplate and incubated for 24 hours. Oxygen consumption rate (OCR) and extracellular acidification rate (ECAR) by cells were measured in XF assay media (unbuffered DMEM containing 2 mmol/L glutamine and 1 mmol/L pyruvate) after subsequent injections of glucose (final = 10 mmol/L), oligomycin [O4876; Sigma (final) = 1 μmol/L], carbonyl cyanide 4-(trifluoromethoxy)phenylhydrazone [FCCP, C2920; Sigma (final) = 1.5 μmol/L], rotenone [R8875; Sigma (final) = 1 μmol/L], and antimycin A [A8674; Sigma (final) = 1 μmol/L] according to the manufacturer’s protocol. After each experiment, live cells were counted by trypan blue exclusion and ECAR and OCR values were normalized, although minimal cell death was observed.

### 4.10 Intracellular Reactive Oxygen Species (ROS) detection assay

To measure the total level of intracellular ROS, cells were stained with 100 μl H2DCFDA (2’,7’ –dichlorofluorescein diacetate) in 1x HBSS 72 hours after being seeded. Cells were washed with 1x PBS twice and diluted in 1mL of 1x HBSS. Fluorescent signals were detected using BD FACSCaliburtm at 488-nm wavelength and data was analyzed by the Flowing 2.5.1 software.

### 4.11 Mitochondrial superoxide detection assay

Cells were grown at 37°C and 5% CO2 for 72 hours and stained with 5μM MitoSOX in 1x HBSS for 10 min at 37°C in dark. Stained cells were washed and diluted in 1mL of 1x HBSS. Fluorescent intensity was determined at 586 nm using BD FACSCaliburtm.

### 4.12 Transmission Electron Microscopy and Morphometric Analysis (TEM)

MDA-MB-231 empty vector and shTRPML1 transduced cells were seeded at 90% confluency 24 hrs before being fixed with 1% glutaraldehyde in 0.1 M sodium phosphate buffer, pH 7.3. The fixation buffer was replaced with a wash buffer containing 100 mM Tris (pH 7.2) and 160 mM sucrose for 30 min. Cells were then washed two more times for 30 min with phosphate buffer (150 mM NaCl, 5 mM KCl, 10 mM Na_3_PO_4_, pH 7.3). After the third wash step, cells were treated with 1% OsO_4_ in 140 mM Na_3_PO_4_ (pH 7.3) for 1 hr. Following OsO_4_ treatment cells were washed two times with H_2_O and stained with saturated uranyl acetate for 1 h, dehydrated in ethanol, and embedded in Epon (Electron Microscopy Sciences, Hatfield, PA). Dehydrated cells were then sectioned (~60 nm) and stained with both uranyl acetate and lead nitrate. The resulting grids were then imaged using a Philips CM-12 electron microscope (FEI; Eindhoven, The Netherlands) paired with a Gatan Erlangshen ES1000W digital camera by a blinded independent observer (Model 785, 4 k 3 2.7 k; Gatan, Pleasanton, CA). Mitochondrial morphology and matrix density were quantified using ImageJ (NIH) software. Mitochondrial circularity was determined using the formula 4π * area/perimeter^2^ where 1 = *perfect circle*. The mitochondrial aspect ratio was determined by dividing the centerline length by the average width. Cross-sectional surface area, mitochondrial matrix density, and percent 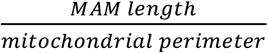 were *mitochondrial perimeter* determined by tracing the outer mitochondrial membrane. Lastly, mitochondrial density was determined by measuring the mean electron density of the mitochondrial matrix after background subtraction.

### 4.13 Reagents

Cell culture media, FBS, PBS, HBSS, and penicillin/streptomycin antibiotic were purchased from Invitrogen/Thermofisher scientific. MTT, oligomycin, FCCP, rotenone, actinomycin and doxorubicin were bought from Sigma-Aldrich.

### 4.14 Statistical analysis

Data are presented as mean ± SEM. Statistical comparison was performed using analysis of variance (ANOVA) and Student’s *t test.* Results presented are mean ± SEM. P values of < 0.05 are considered statistically significant. *: P < 0.05, **: P < 0.01, ***: P < 0.005.

## Funding & Acknowledgement

This work was supported by Dalhousie University Startup funds to YEH, and by the National Heart, Lung, and Blood Institute (NHLBI) grant R35-HL150778 to MT.

## Availability of data and materials

All data and materials used in the current study are available from the corresponding author upon request.

## Consent for publication

Not applicable

